# Exploring light chain cardiotoxicity in AL amyloidosis: Impact on hiPSC-derived Cardiomyocyte Activity

**DOI:** 10.1101/2025.10.23.684089

**Authors:** Calamaio Serena, Frosio Anthony, Melgari Dario, Broggini Luca, Sonzini Federica, Prevostini Rachele, Anastasia Luigi, Pappone Carlo, Nuvolone Mario, Palladini Giovanni, Ricagno Stefano, Rivolta Ilaria

## Abstract

**Aims:** Immunoglobulin light chain (AL) amyloidosis is a protein misfolding disease characterized by the systemic deposition of amyloid fibrils derived from monoclonal light chains (LCs). Cardiac involvement is the major determinant of prognosis and mortality, and beyond fibril accumulation, soluble cardiotoxic LCs play a critical role in disease progression. While current in vivo models like *C. elegans* and murine systems have demonstrated LC toxicity, they lack human relevance or fail to capture soluble LC-induced cardiotoxicity. This study aimed to characterize the electrophysiological effects of cardiotoxic LCs on a human-relevant model using human-induced pluripotent stem cell-derived cardiomyocytes (hiPSC-CMs).

**Methods and Results:** Two amyloidogenic cardiotoxic LCs (H3 and H6) from AL patients and one non-cardiotoxic LC (M10) from a multiple myeloma patient were biophysically characterized and tested in hiPSC-CMs at clinically relevant concentrations. Electrophysiological recordings revealed that H3 and H6 significantly reduced spontaneous action potential (AP) firing frequency and maximal upstroke velocity (dV/dt) in hiPSC-CMs, indicating impaired excitability. H6 also shortened AP duration. H3 exposure led to a ∼40% reduction in peak sodium current density and altered inactivation kinetics of the L-type calcium current, without affecting major pacemaker or repolarizing potassium (I_Kr_ or I_Ks_) currents. In contrast, M10 had no effect on any measured parameter, validating the model’s ability to discriminate toxic from non-toxic LCs.

**Conclusion:** This study demonstrates that hiPSC-CMs provide a clinically relevant human model to investigate LC-induced cardiotoxicity. Cardiotoxic LCs exert distinct but converging electrophysiological impairments, including disruption of sodium and L-type calcium currents, contributing to reduced excitability and altered AP morphology. These findings provide mechanistic insights into AL amyloidosis-related cardiac dysfunction and establish a foundation for future therapeutic screening targeting soluble LC toxicity in a human context.

## 1. Introduction

Immunoglobulin light chain (AL) amyloidosis is a protein misfolding disorder characterized by the conversion of patient-specific immunoglobulin light chains (LCs) from their native state into highly organized amyloid fibrils^1^. The disease originates from the hyperproliferation of a plasma cell clone, leading to overproduction of LCs and their secretion into the bloodstream^2^. The structural details of such amyloids from several patients and from different tissue deposits have recently been reported^3-8^. Clinical manifestations arise from the deposition of fibrillar aggregates in various organs, ultimately causing organ dysfunction^9^. However, cardiac involvement is a hallmark of AL amyloidosis, with over 75% of patients presenting significant cardiac manifestations^10^. This typically appears as a progressive infiltrative cardiomyopathy characterized by electrical abnormalities and advancing heart failure, and it is the main determinant of morbidity and mortality in these patients^9^. Beyond the structural damage caused by LC amyloid deposits in the heart, an additional and crucial pathogenic factor is the direct cardiotoxicity of soluble pre-amyloid LC species^11-16^. Clinical observations strongly suggest a relevant role of circulating toxic LC species, as reducing their concentration through anti-clonal chemo/immunotherapy can lead to rapid improvements in cardiac function and patient outcomes, even in the absence of a measurable decrease in amyloid cardiac deposits^17-18^. Experimental models have further corroborated this concept, showing that cardiotropic amyloidogenic LCs alone, without amyloid fibrils, adversely affect the viability of human and rodent cardiac cells by inducing oxidative stress, disrupting protein homeostasis, and impairing mitochondrial function^16,19-23^. The nematode *C. elegans* is a well-established model to assess LC toxicity *in vivo*. The administration of aliquots of natively folded LCs derived from AL patients with cardiac involvement resulted in severe structural and functional damage to the worm pharynx, the functional analogue of the vertebrate heart, and is closely associated with increased production of reactive oxygen species (ROS) and mitochondrial injury^24,25^. Interestingly, non-amyloidogenic LCs derived from multiple myeloma (MM) patients do not exhibit such significant toxicity in nematodes^24^. The same concept was corroborated by exploiting a recently developed transgenic nematode expressing human amyloidogenic LCs derived from cardiac AL from an MM patient, further expanding the study of the pathophysiological mechanisms of soluble toxicity in AL amyloidosis in *C. elegans*^26^. Martinez-Rivas and colleagues successfully recapitulated the structural damage caused by LC amyloid deposits in a murine model of AL amyloidosis. However, that model lacked the pathogenic profile related to soluble pre-amyloid toxic species since no direct LC cardiotoxicity was observed^27^.

The soluble LC cardiotoxicity correlates well with high fold stability, low flexibility, and conformational variability^15,28,29^ and specific binders of a non-native open conformation of dimeric LC efficiently block LC toxicity *in vivo*^30^. To date, however, the exact nature of the cardiotoxic species has not been fully elucidated. Moreover, although existing models have been invaluable for demonstrating the cardiotropic nature of AL LCs and their role in cardiac dysfunction, their limitations—particularly their distance from the human system—have hindered detailed studies of the underlying molecular mechanisms of cardiotoxicity and have reduced the translational potential of experimental findings.

To address this gap and to gain further insights into the effects of cardiotoxic LCs on a clinically relevant, human cardiac cells, we employed human-induced pluripotent stem cell-derived cardiomyocytes (hiPSC-CMs), a widely used experimental model for studying cardiomyopathies. hiPSC-CMs offer numerous advantages over primary human cardiomyocytes and animal models, including an almost unlimited supply of cells, retention of the human genetic background, and the circumvention of ethical and technical challenges associated with other systems. These features make them particularly valuable for preclinical research and investigations into cardiac physiology and disease mechanisms^31,32^. Although hiPSC-CMs remain immature compared to adult cardiomyocytes - exhibiting fetal-like characteristics such as spontaneous pacemaker activity, a depolarized resting membrane potential, elevated I_f_ current expression^33^, and reduced I_K1_^34^ - they still represent a substantial improvement over classical *in vitro* model.

Specifically, we selected two well-characterized cardiotoxic LCs derived from AL patients with severe cardiac involvement (H3 and H6, derived from germline *IGLV1-44* and *IGLV1-51*, respectively) and one non-amyloidogenic LC from a multiple myeloma patient (M10, derived from germline *IGLV2-14*)^28^. The toxicity of the soluble forms of purified native H3 and H6 has previously been demonstrated in *C. elegans*^15,30^. In this study, hiPSC-CMs were incubated with native H3, H6 and M10 under conditions similar to those used in previous *C. elegans* assays, and their effects on the spontaneous action potentials (APs) were evaluated.

Our findings demonstrated that, in contrast to the non-amyloidogenic LC control, cardiotoxic LCs significantly altered AP dynamics by reducing spontaneous firing frequency and disrupting AP morphology. H3, in particular, impaired hiPSC-CMs excitability, likely by affecting ion currents responsible for the rapid AP upstroke. These results indicate that amyloidogenic LCs exert direct and specific toxic effects on hiPSC-CMs, offering valuable insights into the disease’s underlying pathophysiology.

## 2. Materials and Methods

### 2.1 LCs production and purification

Recombinant full-length immunoglobulin LCs from patients with AL amyloidosis or MM were produced according to protocols described in Oberti *et al*.^28^. Briefly, heterologous proteins, produced in the *E. coli* cytoplasm as inclusion bodies, were retrieved and subjected to a renaturation procedure, followed by purification by means of ion exchange and size exclusion chromatography (SEC).

### 2.2 Analytical size exclusion chromatography

Analytical SEC was performed using a Superdex 200 increase 10/600 column operated at 4 °C by an Akta purifying system. Samples were injected into the column extensively equilibrated in 50 mM Hepes pH 8.0, 150 mM NaCl. Runs were imported in GraphPad Prism 9.0 software (CA, USA) for data normalization, visualization and graph generation.

### 2.3 Mass photometry experiment

Mass photometry experiment was done using a Refeyn OneMP instrument (Oxford, UK). The experiments were performed using microscope coverslips, which were assembled into the flow chamber, and silicone gaskets were positioned on the glass surface for sample loading to hold the sample drops with 4 × 4 wells prior to measurements. Contrast-to-mass calibration was achieved by measuring the contrast of tyroglobuline (660 kDa), beta-amylase (224 kDa, 112 kDa, 56 kDa), and bovine serum albumin (66.5 kDa). Calibration was applied to each sample measurement to calculate the molecular mass of each histogram distribution during analysis. For the experiment, H3 was buffer exchanged to PBS pH 7.4 and diluted to a final concentration of 20 nM prior to sample analysis with 3-fold dilution on buffer droplet to a final concentration of 10 nM. For data acquisition, 10 μl of diluted protein was added to the well and mixed, and movies of 60 s duration with 2800 frames were recorded using Refeyn AcquireMP 2023 R1 software in normal measurement mode with regular image acquisition settings. All mass photometry movies of each measurement were processed and analyzed by Refeyn DiscoverMP v2023 R2 software, and Gaussian curves were fit to each histogram distribution, and the mass (kDa), sigma (kDa) and counts were determined.

### 2.4 Circular dichroism spectroscopy

Circular dichroism experiments were carried out on a J-1500 spectropolarimeter (JASCO Corp., Tokyo, Japan) equipped with a Peltier system for temperature control. All experiments were carried out in 50 mM sodium phosphate pH 8.0. LC concentration was 0.2 mg/mL in a cuvette with a pathlength of 0.1 cm. Spectra were recorded from 260 to 200 nm. For each measurement, three replicates were recorded and averaged to yield the final CD spectrum. Data were imported in GraphPad Prism 9 software (CA, USA) for data normalization, visualization and graph generation.

### 2.5 Thermal unfolding ramps

Fluorescence-based thermal shift experiments were performed using a Tycho NT.6 device (Nanotemper) following the changes in the intrinsic fluorescence detected at both 350 nm and 330 nm. Temperature ramps were performed in 50 mM Hepes, 150 mM NaCl, pH 8.0 from 35 °C to 95 °C. Melting temperature is defined as the temperature at which the folding-to-unfolding transition occurs and is the maximum or minimum of the 350/330 nm ratio curve first derivative. Raw data were imported in GraphPad Prism 9.0 software (CA, USA) for data normalization, visualization and graph generation.

### 2.6 hiPSC culture, *in vitro* cardiac differentiation, and LCs incubation conditions

A hiPSC line from a healthy female donor (Thermo Fisher Scientific, cell line: TMOi001-A) was used and maintained on human Biolaminin 521 LN-coated dishes in TeSR-E8 TM medium (Thermo Fisher Scientific)^35^. Cardiac differentiation was conducted as previously described^36^ using the PSC Cardiomyocytes Differentiation Kit (Thermo Fisher Scientific, Italy) on monolayer cultured on Matrigel® hESC-qualified Matrix (Corning, Corning, NY, USA) dishes. For electrophysiological experiments, hiPSC-CMs were detached on day 21 of differentiation, purified using magnetic beads (Miltenyi Biotec, Germany) according to the manufacturer’s instructions, and replated as grouped or single cells on Matrigel-coated 35mm dishes (VWR, Italy). Cells were allowed to adhere for at least 48 h before conducting experiments. hiPSC-CMs were incubated for 24 h with H3, H6, and M10. LC were diluted in PBS containing Ca^++^ and Mg^++^ (Thermo Fisher) and H3 was used at a concentration of 2.5 µM, or 5 µM to evaluate biocompatibility and at 1 µM, 2.5 µM, or 5 µM to study the electrophysiological impact. These doses were selected based on previous studies^15^ and in line with the concentration present in the patients’ serum^28^. H6 and M10 effects were evaluated only at the concentration of 2.5 µM.

### 2.7 Immunofluorescence staining

For the immunofluorescence staining, hiPSC-CMs at day 21 of differentiation were purified as described and plated on a glass slide coated with Matrigel®. After 24 h, cells were fixed with PFA 4% in PBS for 15 min at room temperature (RT). Staining was performed as previously described^37^. The primary antibody (Anti-Cardiac Troponin T antibody [1C11] mouse monoclonal antibody ab8295, Abcam distributed by Prodotti Gianni s.r.l., Italy, diluted 1:800) was incubated overnight at 4 °C. The secondary antibody (Donkey Anti-mouse Alexa Fluor 594 A-21203, Thermo Fisher Scientific Italia, Italy, diluted 1:600) was incubated for 1 h at RT in the dark. To stain nuclei, 4′,6-diamidino-2-phenylindole (DAPI, Thermo Fisher Scientific Italia, Italy, diluted 1:5000) was used and incubated for 10 minutes at RT in the dark. Images were acquired with a LSM710 confocal microscope (Zeiss, Germany), equipped with a 63× oil immersion objective, as single optical section. Image processing was performed using Zeiss ZEN Microscope Lite software version and ImageJ 1.48V.

### 2.8 Metabolic activity assay (MTT)

hiPSC-CM purified on 21 days were seeded at confluency of 100k cells/cm^2^ in 96-well Matrigel®-coated plates (VWR, Italy). After 48 h, the cells were exposed to culture medium containing H3 (for concentration, refer to paragraph 2.6). Untreated cells and lysed cells (with H_2_O_2_) served as negative and positive controls, respectively. After 24 h of incubation, the medium was replaced and 3-(4.5-dimethylthiazolo-2-yl)-2.5-diphenytetrasolium bromide (MTT, Sigma-Aldrich, USA) was added at a concentration of 0.5 mg/mL in cardiomyocyte maintenance medium and incubated for 3 h at 37 °C in a 5% CO_2_ atmosphere. The resulting formazan crystals were dissolved in a 1:1 solution of EtOH and DMSO, and absorbance was measured at 570 nm and 650 nm using a Varioscan LUX microplate reader (Thermo Fisher Scientific, Italy). The difference in absorbance (570-650 nm) was calculated. As this assay is commonly used as a proxy for cell viability, relative cell viability (%) was determined using untreated cardiomyocytes as reference. Two independent experiments were performed, each with three replicates per condition.

### 2.9 Electrophysiology

All experiments on hiPSC-CMs were performed at 37 °C on a manual-patch clamp set-up equipped with a 700B operational amplifier (Molecular Devices, USA). Patch pipettes were pulled with a P1000 puller (Sutter, USA) to a final resistance of 5-8 MΩ for recording spontaneous APs and to 2-3 MΩ for ionic currents. Spontaneous APs were recorded on small groups of beating cells in whole-cell, current-clamp gap-free configuration. Despite the protocol applied for differentiating cardiomyocytes from hiPSCs being designed to obtain ventricular-like enriched cell cultures, some heterogeneity was observed in the AP recordings, which is not uncommon in this field^38^. Therefore, the ventricular-like APs subpopulation was carefully isolated by applying the method published by Burridge and colleagues to the recorded traces^39^. Briefly, cells were classified as ventricular-like if they had a maximal diastolic potential (MDP) < -50 mV, a maximal upstroke velocity (dV/dt) > 10 mV/ms, an AP amplitude (APA) > 90 mV and a ratio between the AP duration at 90% and at 50% of repolarization (APD90/APD50) < 1.4. Any cell that failed to meet just one of these criteria was excluded from the final pool and subsequent analysis.

Protocols and solutions for APs recording and ionic currents are detailed in the Supplementary Data.

### 2.10 Statistical Analysis

Since the purpose of the biochemical and biophysical analysis of LCs was to evaluate the quality of the purified light chains rather than to perform comparative analyses among the three proteins, no technical replicates were included and no statistical testing (p-values) was carried out. For the metabolic activity assay and for the electrophysiology, results are presented as mean ± SEM, with statistical significance set at P < 0.05. Analyses were performed using unpaired t-tests or One- and Two-Way ANOVA, followed by appropriate post-hoc tests (Fisher for t-tests and Fisher, Bonferroni, or Dunnett for ANOVA). N (number of experiments) and n (number of cells) are detailed in Tables and Figures legends.

## 3. Results

### 3.2 Biophysical Characterization of H3, H6, and M10 LC

Analytical size exclusion chromatography of H3 coupled to mass photometry analysis revealed the homogeneity and dimeric nature of the LC (elution volume of 15.5 mL and molecular weight of 47 kDa, Fig. 1). Similarly, the size exclusion chromatography profiles of H6 and M10 showed a predominant peak at around 15.5 mL, revealing their dimeric nature of these LCs in solution (Fig. 1C). Circular dichroism (CD) spectra of the three purified LCs displayed a distinct negative peak at 218 nm, characteristic of β-sheet enriched proteins (Fig. 1F). Additionally, fluorescence-based thermal unfolding assays demonstrated a single, cooperative folding transition for each protein, consistent with the behavior of globular and compact proteins (Fig. 1E). Together, these biophysical analyses confirm that H3, H6, and M10 were pure, well-folded proteins, meeting the critical quality requirements for downstream applications.

**Figure 1.**
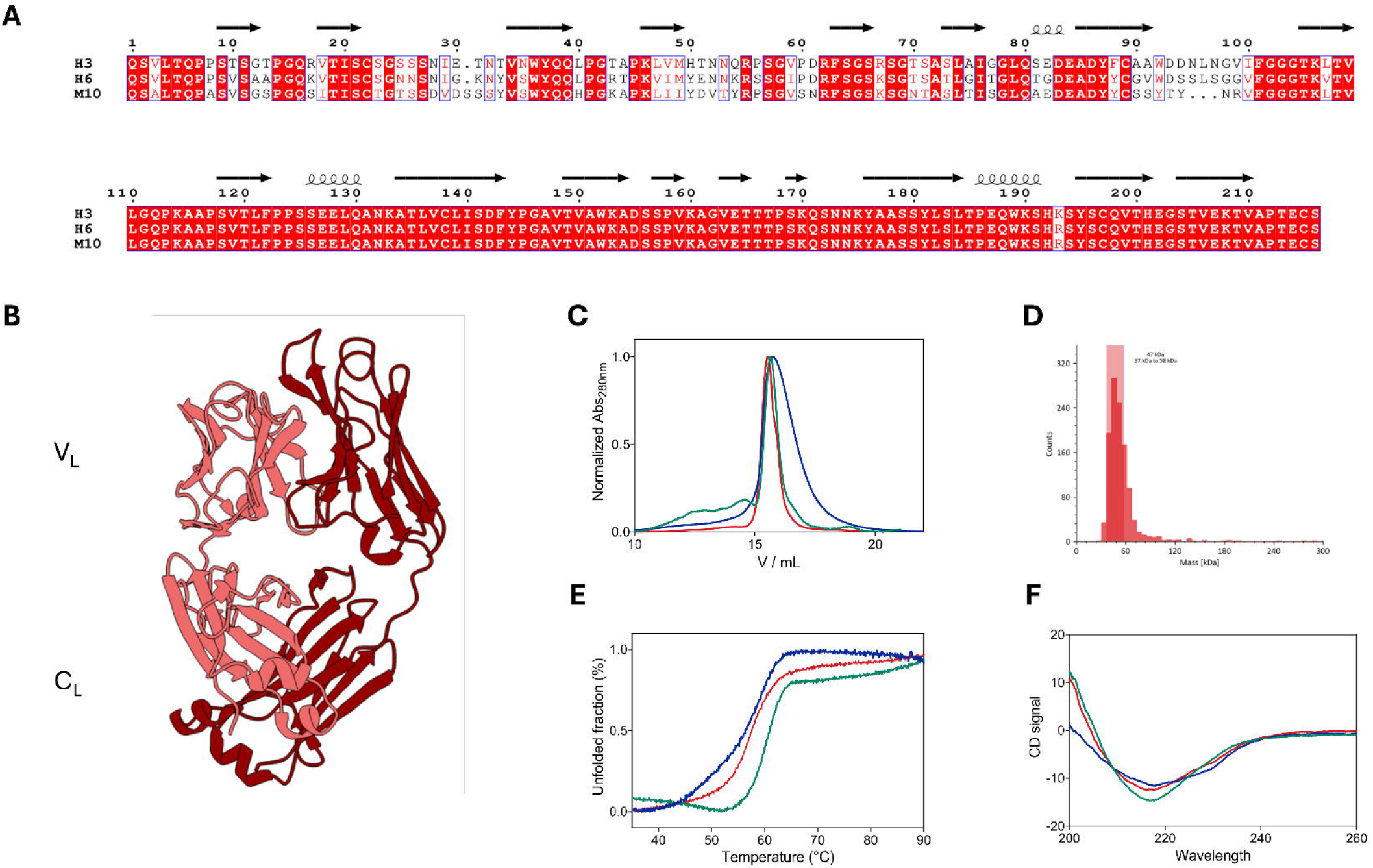
Biochemical and biophysical features of LCs. (A) Alignment of LCs amino acid sequence. (B) Typical immunoglobulin LC dimeric fold with the variable domain (V_L_) V_L_-V_L_ and the constant domain (C_L_) C_L_-C_L_ interfaces stabilizing the homo-dimer. (C) Analytical size exclusion chromatography of LC H3 (red), H6 (blue), and M10 (green) showing a single peak at around 15.5 mL. (D) Mass photometry analysis of LC H3 showing the dimeric nature of the protein. (E) Thermal unfolding ramps of LC H3 (red), H6 (blue), and M10 (green) indicating a single folded-to-unfolded transition. (F) Circular dichroism spectrum of LC H3 (red), H6 (blue), and M10 (green) showing a single negative peak at 218 nm. As stated in the Materials and Methods section, no technical replicates were included and no statistical testing (p-values) was carried out.

### 3.3 Assessment of H3 LC Effects on hiPSC-CMs viability

We assessed whether H3 at the concentration 2.5 µM and 5 µM, representative of concentrations detected in the serum of patients with AL amyloidosis^28^ and previously used in other experimental models^15^, affected hiPSC-CMs viability. To this aim, the MTT assay, an indirect indicator of cell viability, was initially performed on purified hiPSC-CMs (Supplementary Fig. 1). The level of reducing MTT into formazan by mitochondrial enzymes reflects the level of cell metabolism and can thus be considered as a viability index, assuming that only cells with intact metabolic activity (typically viable cells) can convert MTT in formazan. Results indicated that H3 at both tested concentrations did not significantly impact the level of viability (Supplementary Fig. 1B). Based on these findings, we concluded that hiPSC-CMs and the selected LC concentrations could represent a reliable model for further studies.

### 3.3 Cardiotoxic LCs alter spontaneous action potentials in hiPSC-CMs

hiPSC-CMs exhibit a certain degree of heterogeneity that consists in the simultaneous presence of ventricular-like, atrial-like, and sinoatrial node-like APs^39^ Since cardiac AL amyloidosis affects the cardiac conduction system^40^, we focused our study on the impact of the cardiotoxic H3 using spontaneously beating cells with a ventricular-like AP^39^. To provide a comparative analysis, we also tested H6, another cardiotoxic LC, and M10, a non-cardiotoxic LC. hiPSC-CMs were incubated for 24 h with three concentrations of H3 (1 µM, 2.5 µM, and 5 µM), or with the vehicle. H3 significantly reduced the firing frequency of spontaneous ventricular-like APs (Fig. 2A-B for 2.5 µM, and Table 1 for a comparison of 1 µM, 2.5 µM, and 5 µM) at all tested concentrations, in a concentration-dependent manner. Additionally, it markedly decreased the maximal upstroke velocity (dV/dt) (Fig. 2C, Table 1). However, parameters such as maximal diastolic potential (MDP), AP amplitude (APA), and AP durations (APD) measured at 30%, 50%, and 90% of repolarization were unaffected by H3 incubation (Fig. 2D, and Table 1). No changes were induced by the presence of the vehicle alone (data not shown).

**Table 1.** Effect of amyloidogenic light-chains on the parameters of spontaneous APs in hiPS-CMs treated with LC vs untreated CTR (N=number of experiments, n=number of cells, *p<0.05 One-Way ANOVA, Fisher test; # p<0.05 unpaired t-test;)

|  | Frequency (Hz) | MDP (mV) | APA (mV) | dV/dt | APD30 (ms) | APD50 (ms) | APD90 (ms) |
| --- | --- | --- | --- | --- | --- | --- | --- |
| CTR<br>(N=16, n=54) | 2.2 ± 0.1 | -54.8±0.5 | 102.2±0.9 | 32.2±3.2 | 118.0±7.0 | 151.6±8.8 | 183.9±10.5 |
| H3 1 µM<br>(N=5, n=28) | 1.2± 0.1<br>(p= 6.5×10<sup>-7</sup>)* | -56.4±0.9 | 103.1±1.6 | 23.7±2.5<br>(p= 0,046)* | 107.4±7.7 | 142.0±10.3 | 172.7±11.7 |
| H3 2.5 µM<br>(N=6, n=11) | 1.7 ± 0.1<br>(p= 0,049)* | -56.9±1.3 | 104.3±1.8 | 12.3±0.8<br>(p= 0,0012)* | 142.9±18 | 184.28±21.6 | 212.1±23.0 |
| H3 5 µM<br>(N=5, n=18) | 1.8 ± 0.1<br>(p= 0,048)* | -56.1±0.9 | 103.3±1.8 | 14.3±1.0<br>(p=4.03x10<sup>-4</sup>)* | 128.3±13.6 | 161.0±15.7 | 183.1±16.9 |
| H6 2.5 µM<br>(N=5, n=20) | 1.6±0.2<br>(p= 0,028)<sup>#</sup> | -55.6±1.0 | 100.8±1.7 | 21.7±3.5<br>(p=0.08) | 85.4±9.0<br>(p= 0,014)<sup>#</sup> | 112.7±11.2<br>(p= 0,019)<sup>#</sup> | 140.2±13.1<br>(p= 0,027)<sup>#</sup> |
| M10 2.5 µM<br>(N=5, n=31) | 2.5±0.2 | -54.4±0.8 | 100.8±0.9 | 35.7±4.8 | 115.5±9.8 | 144.3±12.1 | 177.6±14.0 |

**Figure 2.**
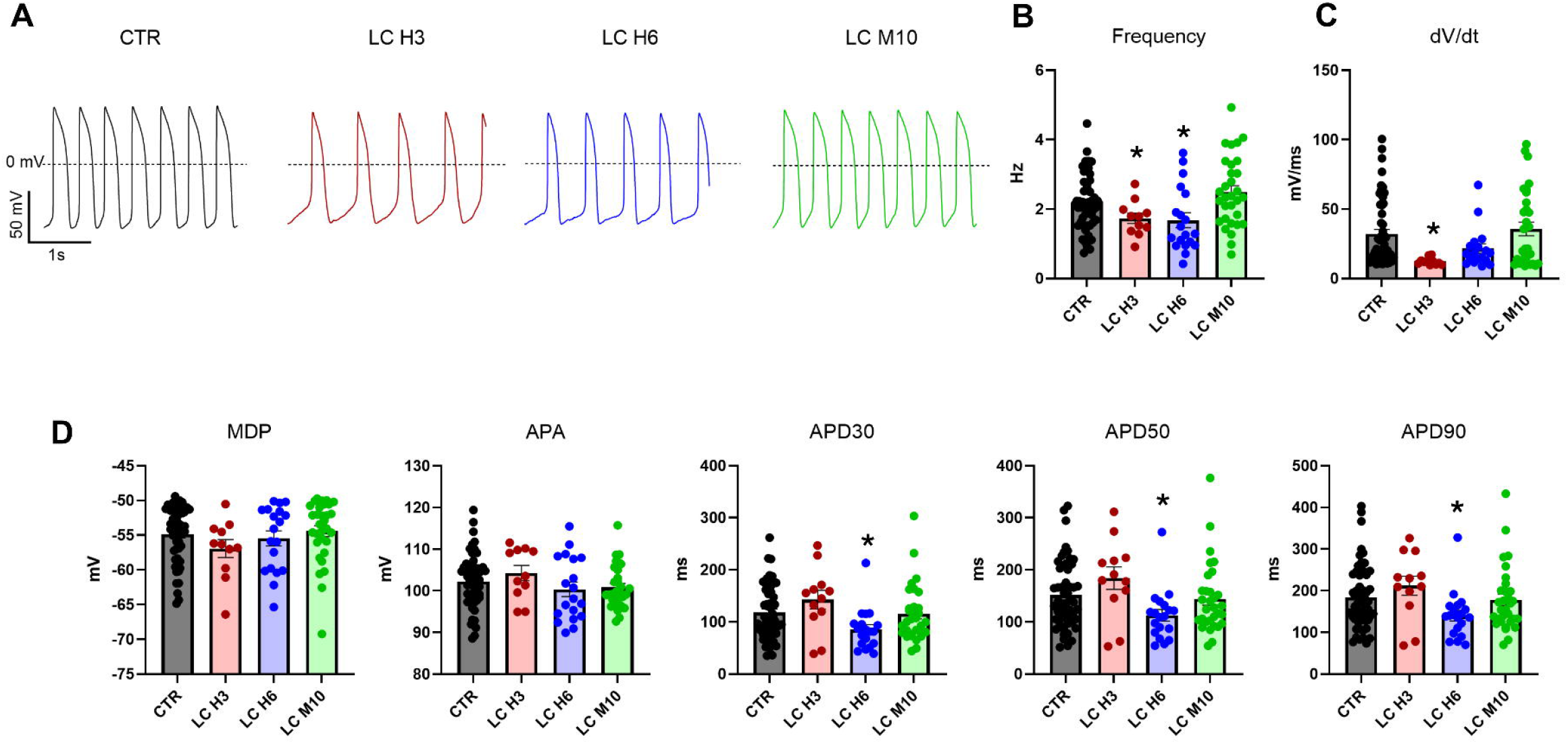
Electrophysiological effects of amyloidogenic and non-amyloidogenic LCs on ventricular-like hiPSC-CMs. (A) Representative traces of spontaneous action potentials (APs) recorded in hiPSC-CMs after 24 h incubation with LC H3 (N=6, n=11), LC H6 (N=5, n=20), LC M10 (N=5, n=31) (2.5 µM each), or vehicle control (N=16, n=54). (B) Quantitative analysis of firing frequency across conditions. LC H3 and LC H6 significantly reduced the firing frequency of spontaneous APs compared to vehicle control, while LC M10 had no effect. (C) Quantitative analysis of the maximal upstroke velocity (dV/dt). Both LC H3 and LC H6 decreased the dV/dt, with LC H6 showing values close to statistical significance, whereas LC M10 exhibited no impact. (D) Analysis of the Minimal Diastolic Potential (MDP), Amplitude of Action Potential (APA), and Action Potential Duration (APD) at 30%, 50%, and 90% of repolarization (APD30, APD50, and APD90). LC H6 exhibited significantly reduced APD at all measured percentages of repolarization, while LC H3 and LC M10 did not alter any of the parameters analyzed. Data are presented as mean ± SEM (N=number of experiments, n=number of cells). Statistical significance was determined using appropriate statistical test, i.e. ANOVA with Fisher’s multiple comparisons test or t-test, \**p* < 0.05 compared to vehicle control. See Table 1 for detailed quantitative values.

The same parameters related to the spontaneous APs were analyzed as key indicators of the impact (if any) of the other selected LCs. Thus, hiPSC-CMs were also incubated with H6 and M10 at the intermediate concentration used for H3 (2.5 µM). Like H3, H6 significantly reduced the spontaneous AP firing frequency of hiPSC-CMs (Fig. 2A-B, and Table 1) as well as the dV/dt, with values being very close to the statistical significance (Fig. 2C, Table 1). Interestingly, H6 also showed a reduced APD at all percentages of repolarization studied (Fig. 2 D, Table 1). By contrast, in hiPSC-CMs incubated with M10 firing frequency, dV/dt, MDP, APA, and APD, closely aligned with the values obtained with the control (Figure 2A-D, and Table 1).

### 3.4 Cardiotoxic H3 affects inward current in hiPS-CMs

The observed impairment of the fast depolarization phase of the APs could suggest a direct effect of H3 on the hiPSCs inward currents (I_Na_ and I_CaL_). To investigate this, the tetrodotoxin (TTX)-sensitive I_Na_ was measured in hiPSC-CMs following a 24 h incubation with H3 at concentrations of 2.5 µM and 5 µM, at which maximal effect was observed. Both concentrations caused a significant ∼40% reduction in the peak current density measured at -10 mV (Figure 3A-B, and Table 2). No significant change in the cell capacitance were observed (22.6 ± 1.3 pF, 20.1 ± 0.7 pF e 20.6 ± 0.9 pF in control, H3 2.5 µM and 5 µM, respectively). A small rightward shift in the voltage dependence of channel activation, significant only at 2.5 µM, was observed (Figure 3C, Table 2), with the magnitude of the shift considered too small to have functional relevance. No significant changes were observed in the fast or slow inactivation time constants at -20 and -10 mV (data not shown), with a general trend towards slowing the process at -30 mV (Table 2).

**Table 2.** Parameters of the ion currents tested in hiPS-CMs exposed to LC H3 (N= number of experiments, n=number of cells, * One or Two-Way ANOVA)

|  | CTR | H3 2.5µM | H3 5µM |
| --- | --- | --- | --- |
| **I<sub>Na</sub> parameters** |  |  |  |
| Current density @-10 mV<br>(pA/pF) | -49.8±6.7 (N=5, n=15) | -29.5±4.2 (N=5, n=14)<br>(p= 9×10<sup>-5</sup>)* | -31.0±5.8 (N=5, n=11)<br>(p=3x10<sup>-4</sup>)* |
| Activation V<sub>1/2</sub> (mV) | -25±0.6 | -22.0±0.8 | -23.9±0.8 |
| Activation slope (mV) | 6.4±0.4 | 6.3±0.5 | 7.1±0.6 |
| Tau fast @ -30mV (ms) | 1.4±0.2 | 2.1±0.6<br>(p=0.02)* | 1.8±0.3 |
| Tau slow @ -30mV (ms) | 8.6±1.7 | 12.6±2.1 | 14.7±3.6 |
| **I<sub>CaL</sub> parameters** |  |  |  |
| Current density @ 0 mV (pA/pF) | -19.1±1.0 (N=5, n=29) | -17.9±1.0 (N=5, n=37) | 19.2±0.9 (N=5, n=51) |
| Activation V<sub>1/2</sub> (mV) | -14.3±0.6 | -12.6±0.4 (p=0.02)* | -13.2±0.4 |
| Activation slope (mV) | 7.1±0.2 | 6.8±0.1 | 6.7±0.1 |
| Tau fast @ -20mV (ms) | 5.2±0.6 | 10.0±1.9 (p=4.3x10<sup>-4</sup>)* | 7.9±0.9 |
| Tau slow @ -20mV (ms) | 52.4±4.6 | 71.1±7.6 (p=0.02)* | 79.4±7.9 (p=3x10<sup>-4</sup>)* |
| **I<sub>f</sub> parameters** |  |  |  |
| Current density @ -125 mV (pA/pF) | -3.1±0.3 (N=4, n=32) | -2.9±0.2 (N=4, n=27) | -3.2±0.4 (N=4, n=25) |
| Activation V<sub>1/2</sub> (mV) | -83.2±1.0 | -80.5±1.1 | -80.7±1.8 |
| Activation slope (mV) | 8.4±0.6 | 6.8±0.5 | 8.2±1.0 |
| **I<sub>Kr</sub> parameters** |  |  |  |
| Current density @ +40 mV (pA/pF) | 0.94±0.08 (N=6, n=23) | 0.86±0.11 (N=6, n=15) | 1.00±0.1 (N=6, n=21) |
| Activation V<sub>1/2</sub> (mV) | -27.7±1.2 | 28.6±4.5 | -28.0±4.3 |
| Activation slope (mV) | 6.6±0.7 | 7.0±1.9 | 5.5±1.9 |
| **I<sub>Ks</sub> parameters** |  |  |  |
| Current density @ +40 mV (pA/pF) | 0.95±0.2 (N=6, n=9) | 1.06±0.18 (N=6, n=9) | 1.53±0.35 (N=6, n=9) |

**Figure 3.**
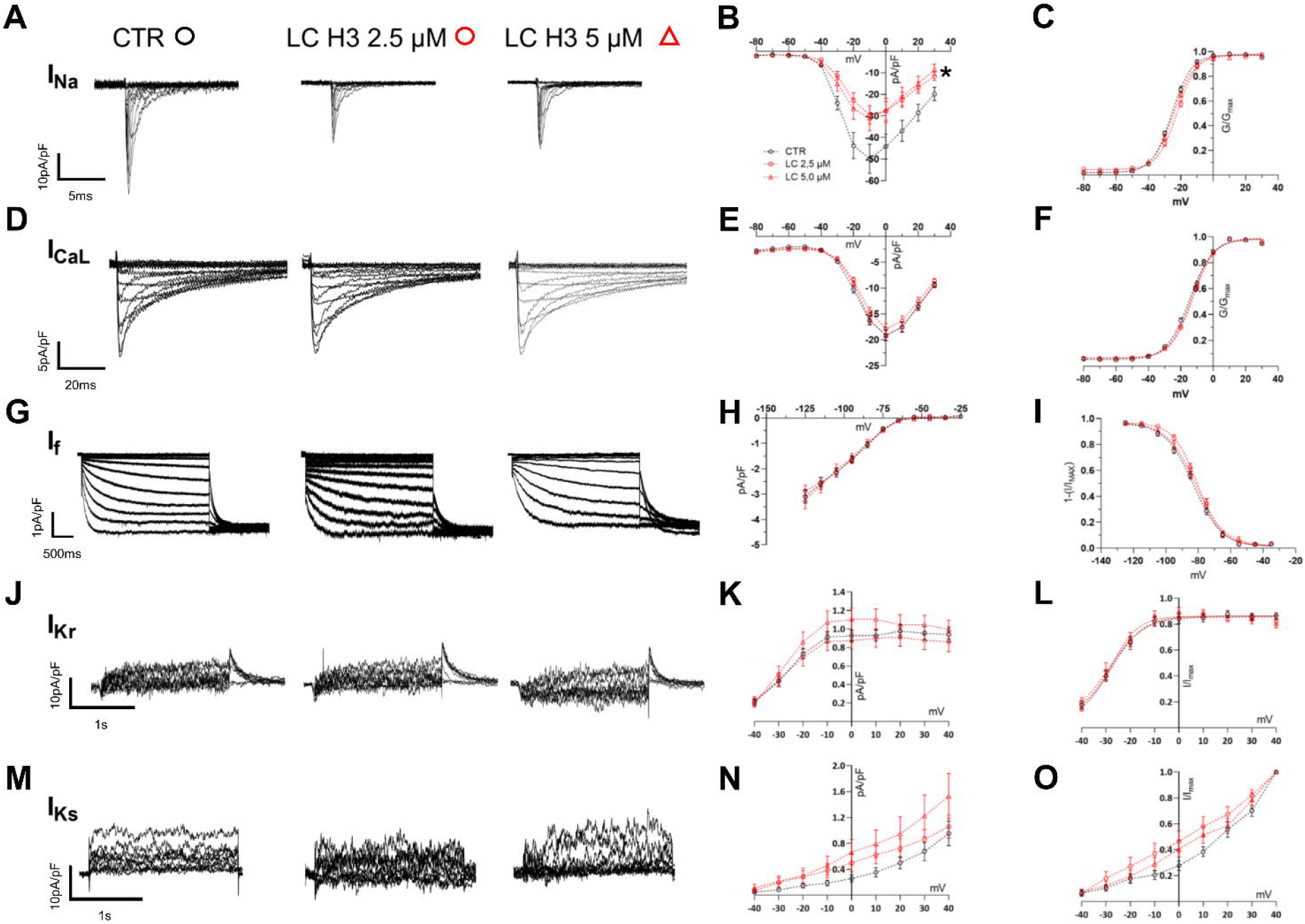
Electrophysiological effects of amyloidogenic LCs H3 (2.5 µM and 5 µM) on ionic currents in hiPSC-CMs after 24 h treatment. (A, B) Sodium current (I_Na_) density showing that LC H3 significantly reduced I_Na_ peak density of about 40% at both concentrations tested. (C) Voltage dependence of I_Na_ activation, showing a minor rightward shift only at 2.5 µM (CTR N=5, n=15; LC H3 2.5 µM N=5, n=14; LC H3 5 µM N=5, n=11). (D, E) L-type calcium current (I_CaL_) density that remained unchanged after incubation with LC H3. (F) A slight rightward shift in voltage dependence of activation observed with LC H3 2.5 µM treatment (CTR N=5, n=29; LC H3 2.5 µM N=5, n=37; LC H3 5 µM N=5, n=51). (G) Pacemaker current (I_f_): neither the amplitude of I_f_ density (H) nor the voltage dependence of channel activation (I) was affected by LC H3 treatment (CTR N=4, n=32; LC H3 2.5 µM N=4, n=27; LC H3 5 µM N=4, n=25). (J) I_Kr_ current: LC H3 at both concentrations did not significantly alter the current density (K) or voltage dependence of activation (L) (CTR N=6, n=23; LC H3 2.5 µM N=6, n=15; LC H3 5 µM N=6, n=21). (M) I_Ks_ current: no change in the current density (N) or in the voltage dependence of activation (O) was observed following the treatment (CTR N=6, n=9; LC H3 2.5 µM N=6, n=9; LC H3 5 µM N=6, n=9). Data are presented as mean ± SEM (N=number of experiments, n=number of cells,). Statistical significance was determined using ANOVA test with Dunnet’s multiple comparisons test, \**p* < 0.05 compared to vehicle control. See Table 2 for detailed quantitative results.

The L-type calcium current (I_CaL_) was isolated as nifedipine-sensitive. Incubation with H3 at 2.5 µM or 5 µM did not affect I_CaL_ peak current density (Figure 3D-E, and Table 2). Similar to I_Na_, the voltage dependence of activation of I_CaL_ was slightly right-shifted with both concentrations, with significance reached only at 2.5 µM (Figure 3F, and Table 2). Interestingly, the incubation with H3 significantly modulated the inactivation kinetic of I_CaL_. When the inactivation decay of the current evoked at -20 mV was fitted with a bi-exponential function, both the calculated fast (τ_FAST_) and slow (τ_SLOW_) time constants were altered following exposure to 2.5 µM LCs. Specifically, τ_FAST_ approximately doubled and τ_SLOW_ significantly increased by 35% compared to control cells. τ_FAST_ also rose substantially at other test voltages (data not shown). At 5 µM LC, τ_SLOW_ was markedly prolonged by 50%, whereas the increase in τ_FAST_ was not statistically significant (Table 2).

### 3.5 Cardiotoxic H3 had no impact on major pacemaker and potassium currents in hiPSC-CMs

To investigate the mechanism underlying the reduced frequency of spontaneous APs observed following incubation with amyloidogenic LCs, the pacemaker current I_f_ was assessed. No significant differences in current density amplitude were found between control cells and those treated with H3 at 2.5 µM or 5 µM (Fig. 3G-H, and Table 2). Likewise, the voltage dependence of I_f_ activation remained unaffected by the treatment (Fig. 3I).

The potential effect of H3 on the two main cardiac repolarizing potassium currents, I_Kr_ and I_Ks_, was also examined. Incubation with either 2.5 µM or 5 µM H3 did not significantly alter current density. (Fig. 3J-K, and Table 2), and the voltage dependence of I_Kr_ activation was unchanged across all conditions (Fig. 3L and Table 2). Similarly, I_Ks_ was not significantly affected by exposure to H3 (Fig. 3 M-O, and Table 2).

### 3.6 Amyloidogenic H6 effect on potassium currents I_Kr_ and I_Ks_ in hiPSC-CMs

Given that H6 at a concentration of 2.5 µM caused a significant shortening of the late repolarization duration in hiPSC-CMs (Fig 2), we tested its effect on I_Kr_ and I_Ks_. Currents were recorded in the same experimental setting employed for LC H3 (see Supplementary data, Materials and Methods section). Incubation of hiPSC-CMs with 2.5 µM H6 did not significantly alter I_Kr_ current density (Supplementary Data Fig. 2A-C). Similarly, I_Ks_ was not significantly affected by the incubation with H6 (Supplementary Data Fig. 2D-F).

## 4. Discussion

Cardiac AL amyloidosis is associated with severe functional consequences, including heart failure with preserved ejection fraction, tachy- and bradyarrhythmias, and various degrees of conduction disturbances. These include atrial fibrillation, ventricular tachycardia, sinus node dysfunction, atrioventricular block, and bundle branch block^40,41^, which are linked to higher rates of ventricular arrhythmias^42^. Notably, these arrhythmias are not necessarily related to morphologic abnormalities or direct amyloid infiltration of the specialized conduction system^43^. To shed light on the cellular mechanisms underlying these clinical manifestations, we investigated the electrophysiological effects of two amyloidogenic and cardiotoxic LCs, namely H3 and H6, on ventricular-like hiPSC-CMs—a powerful model for studying cardiomyopathies that offers critical insights into disease mechanisms while overcoming the limitations of primary human cardiomyocytes. The hiPSC-CM platform employed in this study exhibited electrophysiological properties, including membrane capacitance, upstroke velocity, and resting membrane potential, consistent with those reported for cultures of comparable degree of maturity^31^, thereby supporting their reliability and suitability for detailed electrophysiological analyses.

Here we show that hiPSC-CM AP dynamics and ionic currents display significant alterations upon incubation with cardiotoxic LCs. In particular, the exposure to H3 resulted in a decrease in both the firing frequency of spontaneous APs and the dV/dt, paralleled by a consistent 40% reduction in I_Na_ current density. The absence of major alterations in I_Na_ inactivation kinetics suggests that the primary mechanism of I_Na_ impairment is a decrease in current density rather than changes in gating properties. The parameter dV/dt relates to conduction velocity and serves as an index of sodium conductance in isolated myocytes in phase 0 of the AP. A decrease in dV/dt reflects the presence of diseased cardiomyocytes and identifies a potential arrhythmogenic substrate with an increased risk of arrhythmias^44^. Previous data indicate that there is a strong relationship between dV/dt and diseased myocardium^45^ and traditionally, a decrease in (dV/dt)_max_ and a reduction in I_Na_ have been associated with experimental models of heart failure^46-48^. Decreased dV/dt and a lower frequency of action potentials, another effect of incubation with H3, are also found in progressive cardiac conduction disease^49^, as present in cardiac AL amyloidosis, and may contribute to progressive conduction blocks. However, the observed reduced I_Na_ current density well correlating with the decrease in dV/dt does not rule out the possibility that a reduction in dV/dt may also result from decreased gap junctional coupling and structural changes in the cellular architecture of cardiac tissue, which will be the focus of future studies.

The effects of H3 on the L-type calcium current (I_CaL_) were more nuanced. The prolonged inactivation time constants (τ_SLOW_ and τ_FAST_) observed in the presence of H3 may have an impact on calcium handling^50^, potentially contributing to the overall effect of H3 on heart function and increasing the risk of arrhythmic events.

Surprisingly, the pacemaker current (I_f_), which typically governs spontaneous activity in pacemaker cells, was unaffected by exposure to H3, suggesting that the reduction in spontaneous AP firing frequency is mediated by mechanisms independent of direct modulation of I_f_.

AL is a heterogeneous disease, and the vast variability among LCs, due to genetic rearrangement and somatic hypermutation, results in a unique amino acid sequence for each monoclonal LC^28^. In fact, from a comprehensive perspective provided by the analysis of the APs properties, the effects of incubation with H6 led to the reduction of the spontaneous firing frequency of hiPSC-CMs, as observed with H3 with a less pronounced impact on the dV/dt. However, the reduction in APs duration suggested a potential repercussion on potassium outward currents, in terms of an increase in the outward current density, that, however, was not observed. Thus, at the moment, the mechanism underlying the observed shortening of action potentials thus remains unclear. These results are still consistent with a general remodeling leading to heart failure, but the polymorphic clinical phenotype of cardiac AL may imply that at the cellular level, specific LCs may exert non-identical cardiotoxic effects on cardiomyocytes.

Interestingly, the non-cardiotoxic M10 had no measurable adverse effects on any of the electrophysiological parameters tested. This confirms the ability of this hiPSC-CMs-based analysis to discriminate between LCs which display toxic and non-toxic phenotypes *in vivo*.

In conclusion, the present study demonstrates that hiPSC-CMs are a suitable system to model LC cardiotoxicity in relevant human cell types. H3 significantly impairs key ionic currents, including the sodium current and the inactivation kinetics of the calcium current, in ventricular-like hiPSC-CMs. These alterations contribute to reduced spontaneous AP firing frequency and disrupted AP morphology, which are hallmarks of cardiac dysfunction in amyloidosis. These data however suggest that distinct cardiotoxic LCs may exert different effects on cardiomyocytes nevertheless leading to similar heart impairment. Future research will explore the mechanistic basis of these effects on a wider set of patient-derived cardiotoxic LCs and evaluate potential therapeutic strategies to restore normal cardiomyocyte function in the context of amyloid heart disease.

## Authors Contribution

S.C., A.F., D.M., L.B., S.R., I.R. Substantial contributions to the conception or design of the work S.C., A.F., D.M., L.B., R.P., F.S. Substantial contributions to the acquisition, analysis, or interpretation of data for the work.

S.C., D.M., A.F., L.B., M.N., S.R., I.R. Drafting the work or reviewing it critically for important intellectual content.

G.P., M.N., L.A., C.P., S.R., I.R. Final approval of the version to be published.

