## Supplementary Data for "Exploring light chain cardiotoxicity in AL amyloidosis: Impact on hiPSC-derived Cardiomyocyte Activity"

**Short title: Exploring Cardiotoxicity in AL Amyloidosis**

**Authors**

Calamaio Serena^1,2*^, Frosio Anthony^2*^, Melgari Dario^2*^, Broggini Luca^2,3^, Sonzini Federica^2,3^, Prevostini Rachele^1,2^, Anastasia Luigi^2,4^, Pappone Carlo^2,4,5^, Nuvolone Mario^6^, Palladini Giovanni^6^, Ricagno Stefano^2,3^, Rivolta Ilaria^1,2^.

**Affiliations**

1. School of Medicine and Surgery, University of Milano – Bicocca, Via Cadore, 48, Monza, Italy
2. Institute of Molecular and Translational Cardiology, IRCCS Policlinico San Donato, San Donato Milanese, Italy.
3. Department of Bioscience, University of Milan, Milan, Italy.
4. Faculty of Medicine and Surgery, Vita-Salute San Raffaele University, Milan, Italy.
5. Arrhythmia and Electrophysiology Department, IRCCS Policlinico San Donato, San Donato Milanese, Milan, Italy
6. Amyloidosis Treatment and Research Center, Fondazione IRCCS Policlinico San Matteo, Università Degli Studi di Pavia, Pavia, Italy.

*Contributed equally.

**Corresponding Author:**

Prof. Ilaria Rivolta

School of Medicine and Surgery

University of Milano - Bicocca

Via Cadore, 48

20900 Monza (MB), Italy

**Supplementary Figures**

B

A

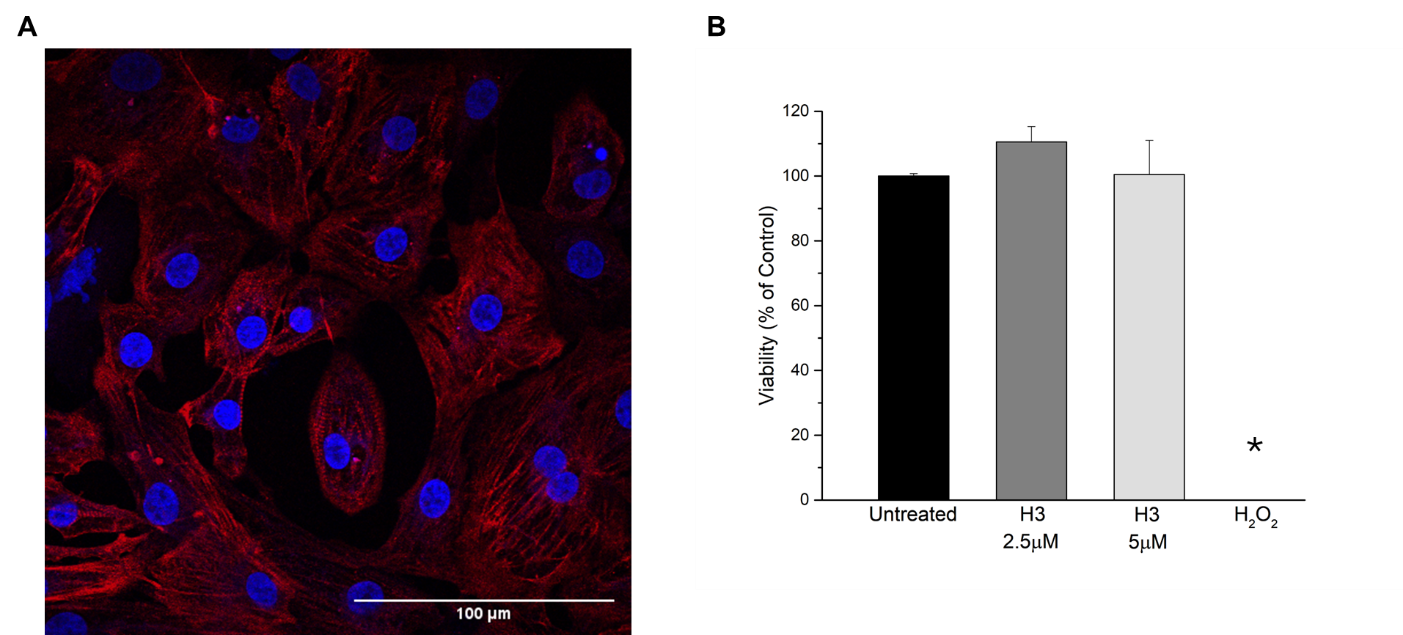

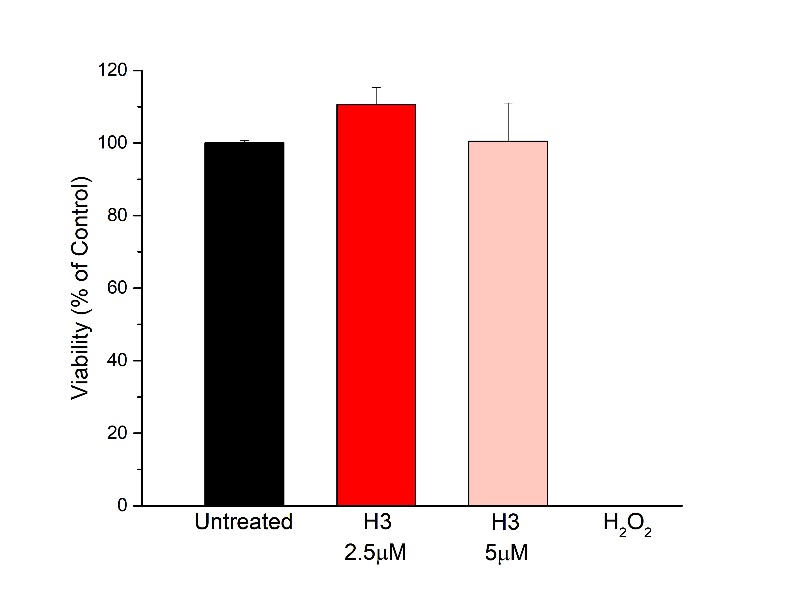

**Supplementary Figure 1. Cellular model and non-cytotoxic assessment of H3 in hiPSC-CMs.** (A) Imunofluorescence of purified hiPSC-CMs at 21 days of differentiation. Confocal image showing the expression of the cardiac marker cTnT (red) and nuclear staining with DAPI (blue). Scale bar = 100 µm. Objective: 63X. (B) Results of the MTT assay confirming that hiPSC-CMs maintained stable viability following 24 h exposure to LC H3 (Untreated 101% ± 10.2; LC H3 2.5 µM 117.6% ±16.8; and LC H3 5 µM 105% ± 2.4; no statistically significant effect was observed), with a significant reduction observed only after H_2_O_2_ incubation (0% ±0.4) used as a negative control to induce cell death. Data are normalized to the mean value obtained for untreated cells and presented as mean ± SEM. Statistical analysis: one-way ANOVA with Fisher’s multiple comparisons test. N=2 technical replicates; n=3 biological replicates.

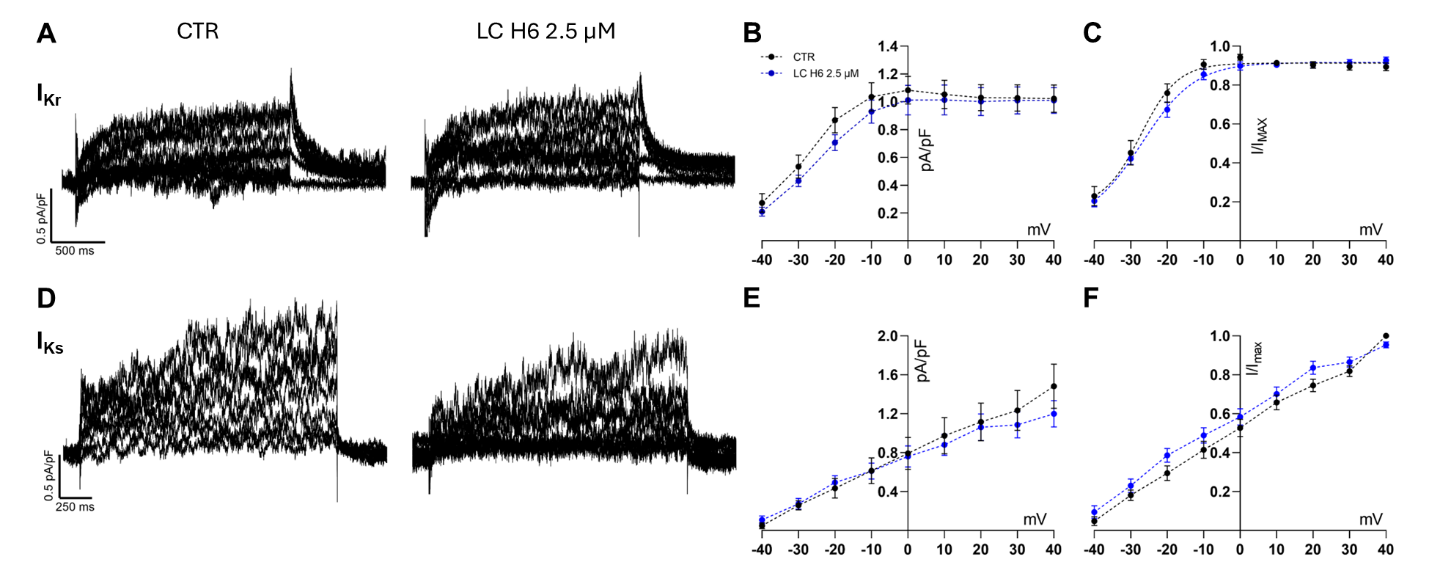

**Supplementary Figure 2**. **Electrophysiological effects of amyloidogenic LC H6 on potassium currents in hiPSC-CMs**. (A) Families of I_Kr_ recorded currents; (B) the current density measured at +40 mV was 1.02±0.09 pA/pF (N=2, n=15) in control cells and 1.01±0.09 pA/pF (N=2, n=23) in presence of 2.5 µM H6. (C) The voltage dependence of I_Kr_ channel activation was unaffected by incubation with LCs, with calculated V_1/2_ values of -26.5±1.5 mV (slope 3.7±0.5 mV), -23.7±1.0 mV (slope 5.6±0.8 mV) for control and H6 incubated cells, respectively. The incubation of hiPSC-CMs with LC H6 2.5 µM did not significantly alter I_Kr_. (D) Families of I_Ks_ recorded currents; the current density (E) was not significantly affected by the incubation with amyloidogenic H6. When measured at +40 mV its amplitude was 1.5 ± 0.2 pA/pF (N=2, n=13) in control cells, and 1.2 ± 0.1 pA/pF (N=2, n=20) after incubation with 2.5 µM H6; (F) the voltage dependence of activation was not altered by the treatment. Data are presented as mean ± SEM. Statistical significance (absent when compared to vehicle control) was determined using 2-Way ANOVA test followed by Dunnet’s multiple comparisons test. ns p > 0.05.

**Supplementary Methods**

**Electrophysiology**

All ionic currents were recorded on single cells in whole-cell voltage-clamp configuration. Sodium and calcium currents were elicited by the same voltage protocol to allow pharmacological subtraction: 10 mV consequential 150 ms steps ranging from the holding potential of -80 mV up to +60 mV. The two currents were pharmacologically separated with 30 µM TTX and10 µM nifedipine. Both currents were sampled at 50 kHz and low-pass filtered at 10 kHz. Steady-state activation and availability curves were fitted with a Boltzmann function: y = 1/(1 + exp((V − V_1/2_)/k)), where y was the relative conductance (G/Gmax) for activation, relative current (I/Imax) for availability, V is the membrane potential, V_1/2_ is the half-maximal voltage, and k is the slope factor. The time course of inactivation of the sodium current was assessed by fitting the decay of inactivating currents measured at -20 mV with a bi-exponential function. The hyperpolarization-activated current I_f_ was elicited with the following voltage protocol: from a holding potential of -35 mV, 3 s 10 mV steps ranging from -25 to -125 mV, followed by a 1500 ms step to -125 mV and a final 400 ms step to 0 mV. I_f_ was sampled at 10 kHz and low-pass filtered at 800 Hz. The repolarizing potassium currents I_Kr_ and I_Ks_ were elicited by the same voltage protocol: from a holding potential of -60 mV, a ladder of 10 mV incremental 2 s steps ranging from -40 to +40 mV was applied before returning to the holding level. I_Kr_ and I_Ks_ were pharmacologically isolated with 3 µM E-4031 and 10 µM chromanol 293B respectively. In all voltage-clamp experiments, cell membrane capacitance and 70% series resistance were compensated. Current density of each cell was calculated by dividing the current amplitude (pA) by the cell capacitance (pF).

Spontaneous APs and all currents were recorded with the same extracellular solution containing (in mM): NaCl 140, KCl 5.4, CaCl_2_ 1.8, MgCl_2_ 10, HEPES 5, Glucose 10 (pH 7.4 with NaOH, 308 mOsm). To record I_f_, 1 mM BaCl_2_ and 2 mM MnCl_2_ were added to the solution. Intracellular pipette solution was the same for APs and I_f_ and potassium currents, containing (in mM): NaCl 10, KCl 120, CaCl_2_ 2, MgCl_2_ 2, HEPES 10, EGTA-KOH 5, Na_2_-ATP 2, Na_2_-GTP 0.1, Na_2_-Creatine Phosphate 5 (pH 7.2 with KOH, 298 mOsm). For calcium and sodium currents, a cesium-based intracellular pipette solution was used with the following composition (in mM): NaCl 10, CsCl 135, CaCl_2_ 2, HEPES 10, EGTA 5, Mg-ATP 2, TEA-Cl 2 (pH 7.2 with CsOH, 298 mOsm).
